# Engineering Particle-based Materials for Vasculogenesis

**DOI:** 10.1101/2023.03.15.532817

**Authors:** Natasha L. Claxton, Melissa A. Luse, Brant E. Isakson, Christopher B. Highley

## Abstract

Vascular networks are critical to the survival of cells within materials designed for regenerative medicine. Developing approaches to vascularize three-dimensional (3D) *in vitro* models that recreate tissue physiology and 3D tissue constructs for regenerative medicine remain an important focus of tissue engineering. Granular hydrogels are emerging as a promising class of materials for the regeneration of damaged tissues and fabricating tissue constructs. While granular hydrogels have supported vasculature formed by angiogenesis and fabrication processes that establish channels, parameters for designing these materials to support formation of vasculature by vasculogenesis from cells contained within these materials are not fully understood and remain largely unexplored. In this study, vasculogenesis within 3D granular hydrogels formed from polyethylene glycol (PEG) microgels are studied for its potential to establish a microvascular network within this class of materials. Self-organization of endothelial cells into networks within hours is observed in the presence of fibroblasts, and the effects of cell adhesive ligands (RGD) and porosity are measured. Increasing porosity is observed to enhance vasculogenesis while the addition of RGD impairs microvessel network formation. This work establishes parameters that support robust microvasculature formation within granular hydrogels that might be broadly applicable to this class of materials, with implications for other morphogenetic processes in 3D systems.

## I. Introduction

Three-dimensional (3D) tissue constructs, and *in vitro* models are needed to recapitulate important characteristics of tissue physiology and disease that cannot be modeled in 2D and to engineer tissue replacement ^[1]^. 3D tissue structures that recreate complex compositions of cells and extracellular environments enable platforms for drug discovery and aid in regenerative medicine therapies. Within 3D tissue constructs, the development of vascular structures is critical to recapitulating the complexity of tissue physiology that can be studied. Recreating the interactions between vasculature and surrounding tissue ^[2]^ and its absence is detrimental to construct growth and survival. However, it remains challenging to vascularize 3D constructs as well as 3D organoid and cell spheroid cultures ^[3]^.

Hydrogels in which vasculogenesis has been seen include those derived from natural as well as synthetic materials. Robust 3D microvascular networks can form in fibrin ^[4]^ hydrogels without chemical modification of the polymeric backbone. In many advanced hydrogel biomaterials used for tissue engineering ^[5,6]^, carefully designed chemical modifications impart biochemical and biophysical functionalities and are used for crosslinking. In these engineered materials, hydrogel network structure can be designed to permit dynamic cellular activity, such as vascular morphogenesis, within the hydrogel bulk. These “permissive” materials include enzymatically degradable hydrogels that allow cell mobility and microvessel formation in 3D ^[7–9]^, porous hydrogels, and hydrogels crosslinked with dynamic chemical bonds ^[10,11]^ that allow network rearrangement as cells proliferate and move ^[12]^.

Particle-based (or granular) formulations of hydrogels ^[13,14]^ have the potential to impart upon hydrogel properties that render them permissive to 3D cellular activity without engineering the hydrogel network to yield or degrade. Granular hydrogels’ bulks consist of hydrogel microparticles, with controlled space between them ranging from completely jammed to highly porous^14^. Designed porosity ^[15]^ and ability of microgels that comprise the bulk of a granular gel to move under stress ^[13,16–18]^ inherently offer potential for cellular movement and tissue morphogenesis without the use of degradable or dynamic chemistries in the design of the hydrogel’s crosslinking or polymer backbone. Seminal work with microporous annealed particle (MAP) gels demonstrated the great potential that designed porosity offered in promoting cell infiltration and regeneration of vascularized tissue ^[19]^. At the same time, the liquid-like properties of granular hydrogels were also demonstrated to enable cells, including vascular endothelial cells, to be suspended within hydrogel particles ^[20]^ within an established environment that permitted cell migration and proliferation ^[16]^. Recent work has continued to demonstrate the broad potential of granular materials in regenerative medicine and tissue engineering. For example, in new injectable and extrudable materials ^[14,21]^, in the development of materials for biofabrication and 3D printing ^[6,9,11,22]^, and in diverse processing of hydrogels into granular formulations ^[19,21]^.

To approach the broad challenge of vascularizing engineered tissue constructs, promoting vasculogenesis from endothelial cells included within granular formulations of hydrogels would enable vascularization of diverse hydrogel chemistries. Porosity, which has been shown to strongly support vascular ingrowth ^[19,22]^, and viscoelastic properties that influence processes—including vasculogenesis within continuous (non-granular) hydrogels ^[10]^— might be controlled in new ways in granular materials. Granular materials can relax under applied stresses that are above a yield stress ^[10,23]^, a property that stands out in granular materials in which interparticle crosslinking ^[13,24]^ (or annealing) ^[19]^ has not been established. Stress relaxation of the extracellular environment has been identified as an important feature in governing cell behaviors in 3D environments, including cell migration, morphogenesis of cellular features and tissue structures ^[25]^, and vasculogenesis ^[10,26,27]^. The design of hydrogels with stress relaxation characteristics like biological tissues ^[23,28]^—a challenge using traditional bulk hydrogel approaches—might be addressed through granular hydrogel formulations. Permissive materials with designed porosity and yielding, may generally enable the morphogenesis of tissue structures from cell populations embedded within them. In the specific and important challenge of vascularizing hydrogels, permissive formulations of hydrogels may also allow vasculogenesis from endothelial cells and support cells within diverse hydrogel materials.

In this work, we have studied vasculogenesis in 3D granular hydrogels formed from polyethylene glycol (PEG) microgels, whose internal—or intraparticle—crosslinking is covalent, but where the bulk, granular hydrogel has no interparticle crosslinking. Our work relates porosity and the presence of cell-adhesive ligands within granular hydrogel formulation to the emergence of a microvascular network, and we observed an extensive microvasculature network in a co-culture of endothelial cells and fibroblasts. Microvascular network formation in granular hydrogels occurred rapidly, over hours, without PEG degradation, offering an alternative to approaches that have pioneered proteolytically active hydrogel chemistries ^[29]^ and dynamic crosslinking ^[10]^.

## II. Results and Discussion

Bioactivity can be engineered within a PEG hydrogel through selective functionalization of the hydrogel network. PEG hydrogels have been designed to support 3D tissue engineering, vasculogenesis ^[30]^, and organoid culture ^[31]^. To vascularize constructs or enable the morphogenesis of tissue structures in a 3D PEG hydrogel, permissive chemistries are needed within the hydrogel network, typically at sites of macromer crosslinking. Here, we use a non-degradable PEG formulation to form microgel particles that are combined into a granular material with no interparticle crosslinking. As particle-based hydrogels have inherent porosity and stress-yielding properties, they have served as a class of materials for studying dynamic cell behaviors ^[13,26,32]^. With research in the field of biomaterials exploring the potential of these materials as tissue engineering scaffolds, we looked to understand whether they would support vasculogenesis as a method to establish vasculature that is critical to engineered tissues. Vasculogenesis is a dynamic process which has been demonstrated in bulk hydrogels engineered to degrade ^[30]^, contain pores ^[33]^, or undergo stress relaxation ^[10]^. Here, we introduced endothelial and support cells into a granular PEG hydrogel, an inherently dynamic biomaterial system, to understand how they might be designed to support vasculogenesis (Figure 1A).

**Figure 1:**
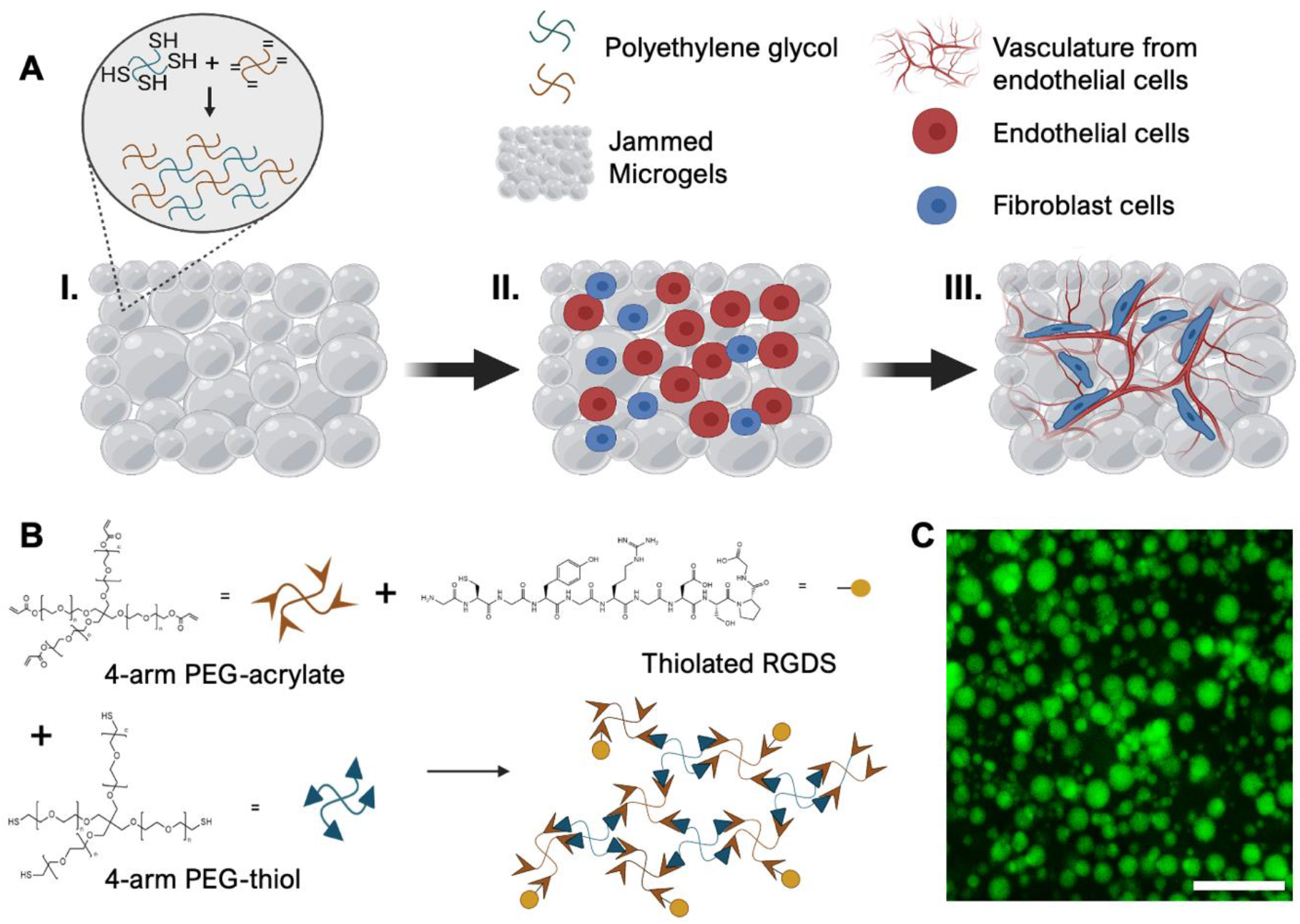
A granular hydrogel is formed from many individual PEG microgels, where excess fluid has been removed from between the microgels to result in jamming. **A**. Schematic illustration of (***I***) a granular hydrogel formed from jammed microgels (gray), where individual microgels are formed in emulsion through the crosslinking of polyethylene glycol (PEG)-acrylate and PEG-thiol (inset: hydrogel network schematic); (***II***) by including endothelial cells, a vascularized construct can be formed using endothelial cells (red) and fibroblasts (purple); (***III***) over time, cells mature into a microvascular network formed from endothelial cells supported by fibroblastsw. **B**. PEG macromer and peptide chemistry, as well as hydrogel network schematic: 4-arm PEGs are combined, with acrylates reacting with thiols at elevated pH. Stoichiometric control can leave unreacted acrylates for further functionalization. **C**. Representative image of microgels (2 MDa FITC-dextran is encapsulated for visualization and size characterization) created by emulsification, dispersed in excess water before jamming (scalebar = 200 μm).

PEG microgels were produced via emulsification of PEG-thiol and PEG-acrylate in solution (Figure 1B), with aqueous droplets crosslinking to form microgels via Michael-addition while in emulsion (Figure 1A, I). Microgels could easily be modified with cysteine-terminated peptides added to the solution, such as the fibronectin-derived cell adhesive ligand RGD used here (Figure 1B). To visualize PEG microgels, high molecular weight (2 MDa) fluorescein-isothiocyanate-dextran was entrapped within the PEG during crosslinking. After recovery from oil, washing to remove surfactants, and incubation in excess water prior to being used to create granular hydrogels, these particles (Figure 1C) were found to have a size distribution of mostly <40 μm average diameter with 21-40 μm diameter constitution the majority at ∼30% (Figure 2A). Microgels suspended in water were centrifuged at 3,095 rcf (x *gravity*) for five minutes, and the resulting granular hydrogel formed from jammed microgels was visualized as having densely packed particles (Figure 2B). The granular bulk hydrogel formed through the concentration of PEG microgels (Figure 2C, I) is stabilized by the physical forces of the jammed structure, and maintains its shape after inversion (Figure 2C, II). Towards understanding the mechanical environment cells embedded within these PEG-based granular hydrogels would experience, we characterized the collective properties of the microgels as a granular material (i.e., the bulk properties of the granular hydrogel). When conducting rheological measurements, we found that these granular hydrogels exhibited a storage and loss moduli of 420 Pa and 18 Pa, respectively. In response to increasing strains on the granular hydrogel, the microgels exhibit a solid-like behavior that switches to fluid-like flow at high strains, above a yield stress. This behavior is characteristic of granular hydrogels in which physical jamming interactions, but not interparticle crosslinking, stabilize the bulk materials (Figure 3A). These mechanical properties were within the range typically employed in hydrogels used for soft tissue engineering ^[34]^.

**Figure 2:**
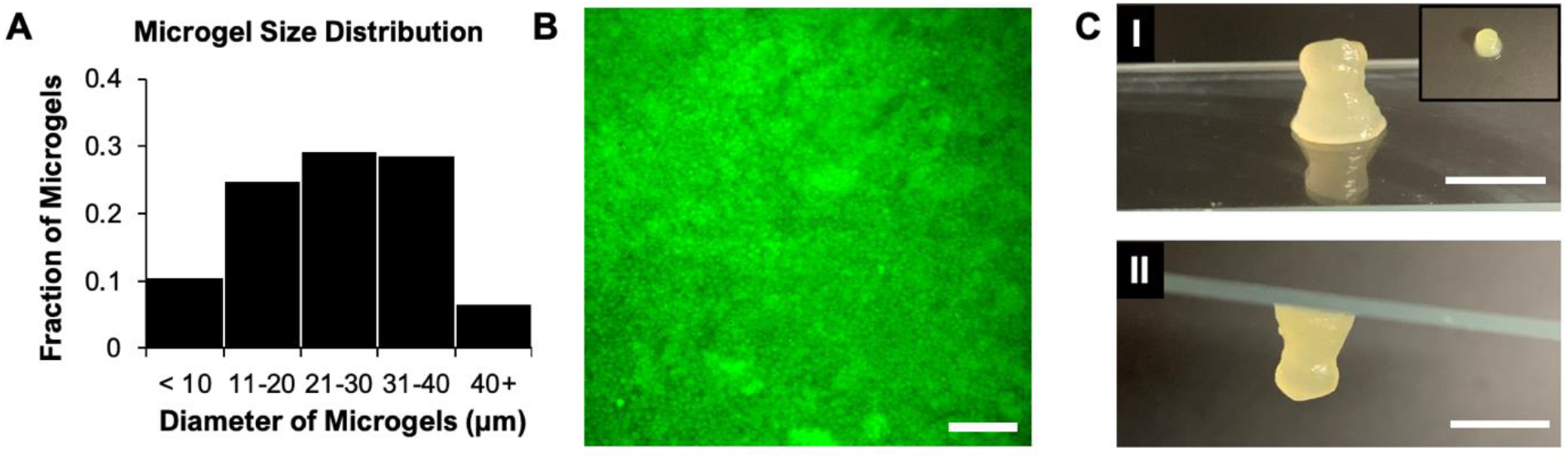
Microgel characteristics and images of a granular hydrogel (microscopic and macroscopic). **A**. Representative size distribution for individual microgels within a granular hydrogel, with average diameter of 30 μm. **B**. A microscopic image of a granular hydrogel formed by jamming of a batch of microgels (containing FITC-dextran) shows concentrated individual microgels (scalebar = 200 μm). **C**. Macroscopic images of a granular hydrogel (***I***) showing the appearance of the granular hydrogel on a glass slide (inset: top view) and (***II***) illustrating its stability even after it is inverted (scalebars = 0.5 cm). The stability of the bulk gel is due to the physical forces of jamming. Here, there is no crosslinking between particles and they will rearrange and flow when stresses exceed the granular hydrogel’s yield stress.

**Figure 3:**
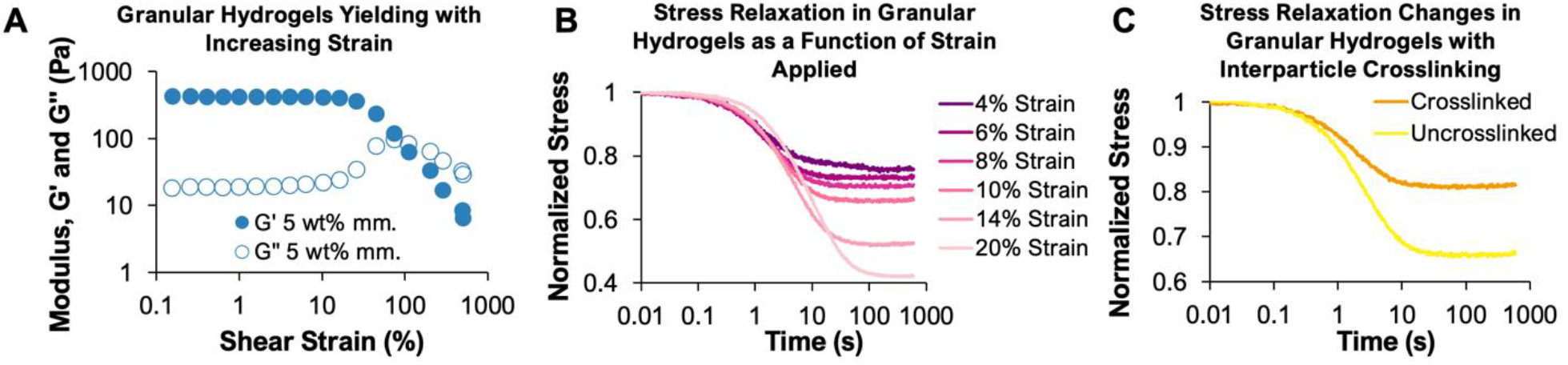
Characterization of rheological and mechanical properties of granular hydrogels formed from PEG microgels. **A**. An oscillatory strain sweep of a 5 wt% PEG-acrylate/PEG-thiol granular hydrogel exhibits elastic behavior below a strain at which the yield stress is exceeded, resulting in fluid-like behavior. **B**. Stress relaxation of 5 wt% granular hydrogels after applying various strains. The granular hydrogels exhibit increasing stress relaxation times as strain increases, with final stress plateauing at lower percent of the maximum stress. **C**. Stress relaxation in granular hydrogels varies as a function of forming crosslinks between particles within the granular material. Interparticle crosslinking decreases the gels’ ability to relax, resulting in higher stresses in the hydrogels after relaxation compared to uncrosslinked granular materials.

Stress relaxation within biomaterials is important for cellular processes that include cell migration ^[23]^, cellular level ^[25]^ and tissue morphogenesis ^[31]^, and vasculogenesis ^[10]^. To quantify the inherent dynamic properties of the granular hydrogels, we used rheology to measure yield stress and stress relaxation in our most jammed formulation. Under increased strains, the granular hydrogel exhibited increasing stress relaxation. The final yield stress (1,900 Pa) exhibited by the microgels after a 20% strain applied plateaus at a lower percent of the maximum stress (Figure 3B). Next, we wanted to examine how interparticle crosslinking would affect mechanical properties in the granular hydrogel. Upon applying a 10% strain to both the uncrosslinked and crosslinked granular hydrogels we saw a decreased ability to relax stress in granular materials where there was particle-to-particle crosslinking (Figure 3C), resulting in higher sustained stresses after relaxation in crosslinked granular hydrogels compared to uncrosslinked granular hydrogels where particle flow was restricted solely by physical jamming interactions.

Porosity in granular hydrogels has been observed to facilitate vascularization of these materials through enhancing cellular ingrowth, both in vivo ^[19]^ and in vitro ^[35]^. Here, we studied how properties of granular hydrogels, including porosity, affect vasculogenesis from cells incorporated into the granular hydrogels during granular hydrogel formation. Rapid establishment of microvascular networks within these materials via vasculogenesis would support applications in tissue engineering, biological studies, and regenerative medicine. We used human umbilical vein endothelial cells and fibroblasts mixed at a ratio 5:1, respectively (Figure 1A, II), and incorporated among PEG microgels during granular hydrogel formation. We monitored microvascular network formation of this co-culture within granular hydrogels (Figure 1A, III) as a function of granular hydrogel design, and compared microvascular network morphogenesis in various 2D and 3D conditions. In cellular experiments in which we wanted to engineer cell adhesion to PEG, RGD was used.

For cellular experiments to initiate vasculogenesis, PEG microgels with and without RGD were homogenously mixed with human umbilical vein endothelial cells (HUVECs) and fibroblasts in a microcentrifuge tube (Figure 4A), then pipetted into a custom polydimethylsiloxane (PDMS) microfluidic device for coculture (Figure 4B). Cell survival 24 hours post seeding was monitored using a calcein AM/ethidium homodimer Live/Dead assay. Cell viability within the granular material was greater than 96% 24 hours after seeding into PDMS devices (Figure 4C). To observe the self-organization of cells into networks of microvascular cells, cocultures were stained with F-actin and DAPI. The emergence of microvascular networks was observed on 2D hydrogel surfaces as well as within hydrogel systems in 3D (Figures 5 and 6). On the surface of a 2D continuous (not granular) PEG hydrogel, the inclusion of RGD and of fibroblasts in coculture with HUVECs were necessary to drive network formation (Figure 5A). We observed that on a 2D PEG hydrogel with RGD, when HUVECs are cultured alone, they form cobblestone-like structure on the hydrogel surface with RGD, reminiscent of their roles in creating a monolayer lining the vascular wall (Figure 5A, I). The introduction of a supporting cell type, fibroblasts, induced formation microvessel-like structures (Figure 5A, II), while not including RGD prevented network formation and resulted in cell aggregation in co-cultures (Figure 5A, III) when cells were found on surfaces. Next, we wanted to test if the addition of growth factors (100 ng/mL human angiopotetin-1 protein and 50 ng/mL VEGF-165A) to the culture medium enhanced vessel connectivity and branching. After four days in 2D culture on a continuous PEG hydrogel with RGD surface, increased and thinner microvasculature formation within the co-culture environment is observed and (Figure 5B) compared to medium without supplemented growth factors (Figure 5A, II).

**Figure 4:**
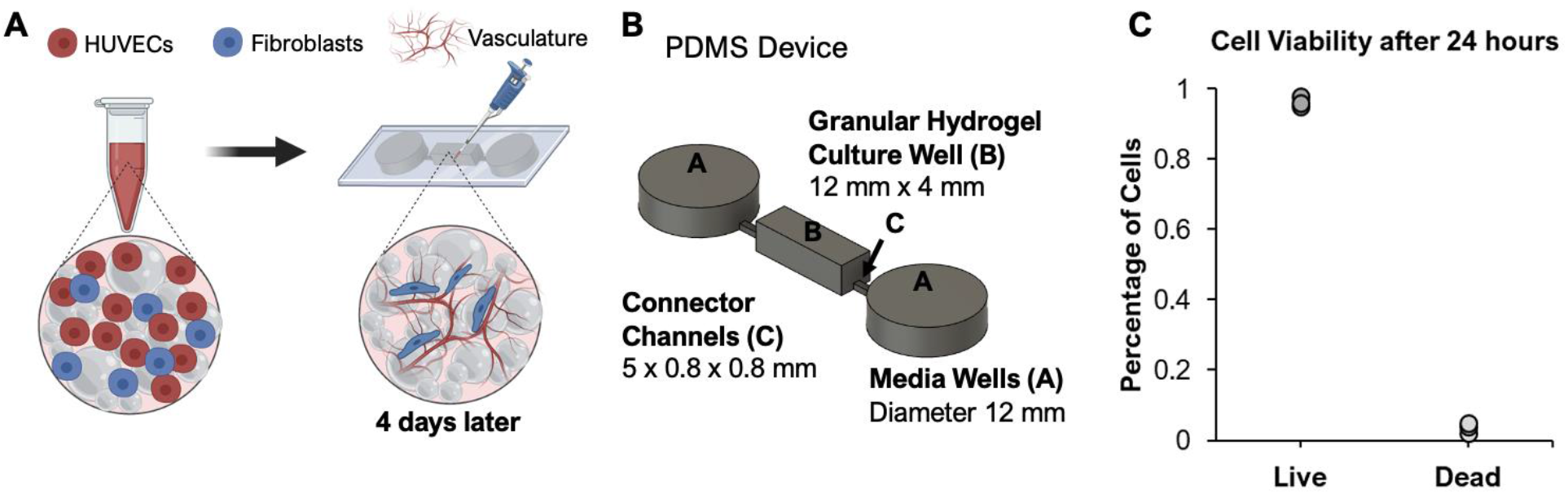
3D culture experiments were performed in PDMS microdevices designed to hold the granular hydrogels in place, absent interparticle crosslinking. **A**. Schematic of co-culture experiment with microgels. HUVECs and fibroblasts were mixed with PEG microgels to form a homogenous mixture, which was then pipetted into a center culture well in the PDMS device. Medium was applied from two circular wells via connector channels. **B**. The 3D computer design of the PDMS mold includes two cell medium wells with 12 mm diameters, a rectangular central well for granular hydrogel culture, and two connector channels for medium perfusion and nutrient/waste exchange. **C**. Cell viability was quantified after 24 hours: >96% viability was observed.

**Figure 5:**
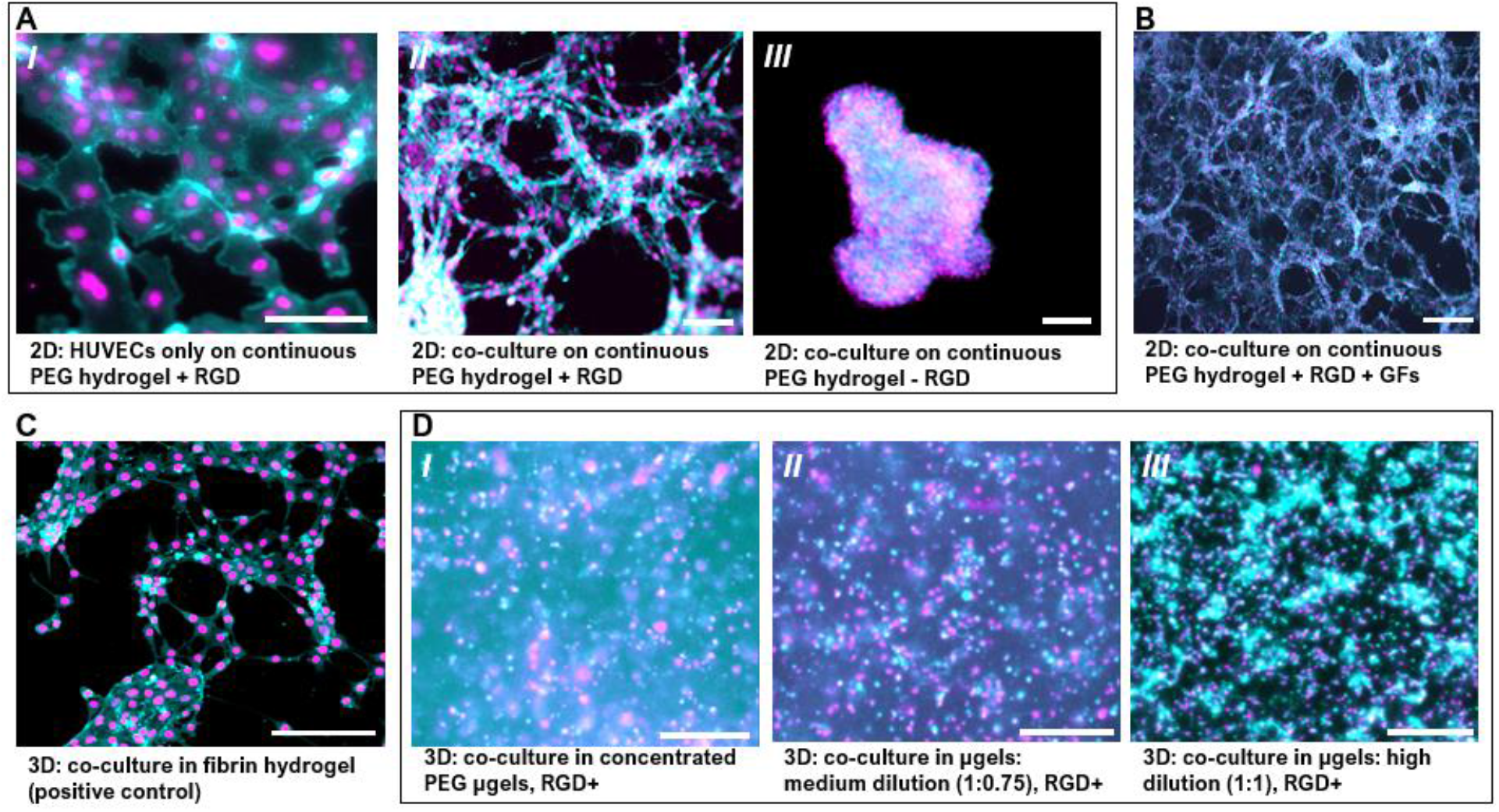
The emergence of vascular structures was observed on 2D PEG hydrogels, a 3D fibrin hydrogel (as a control), and in 3D granular hydrogels formed using PEG-based microgels. F-actin staining (cyan) and nuclear staining (magenta) are shown. **A**. Brightfield images of (***I***) HUVEC monoculture on the surface (2D) of a continuous PEG hydrogel where RGD is added (+RGD) exhibits cobblestone-like cell morphologies reminiscent of blood vessels’ endothelial lining. (***II***) Co-culture of HUVECs with 3T3 fibroblasts on the surface of a continuous PEG hydrogel (+RGD) results in the organization of endothelial cells into a microvascular-like network. (***III***) In the absence of RGD, cells cluster and no cell morphologies seen in vascular structures are observed. **B**. The addition of growth factors to the culture medium (+GFs) enhances microvascular-like network formation in co-cultures of HUVECs and fibroblasts on 2D continuous PEG hydrogel (+RGD) surfaces. **C**. A fibrin hydrogel was used as a control to confirm the organization of microvascular network structures in 3D co-culture from HUVECs and fibroblasts that were homogenously dispersed at the start of the experiment, as has been shown in previous work ^[27]^. **D**. In 3D cultures within PEG-based granular hydrogels that include RGD, microvascular network formation is seen to be enhanced with increasing dilutions of the microgels in the granular hydrogel, from (***I***) fully jammed, to (***II***) a 1:0.75 dilution of the microgels (combining 1 part jammed μgels with 0.75 parts medium), to (***III***) a 1:1 dilution of microgels with cell medium. All scalebars = 200 μm.

**Figure 6:**
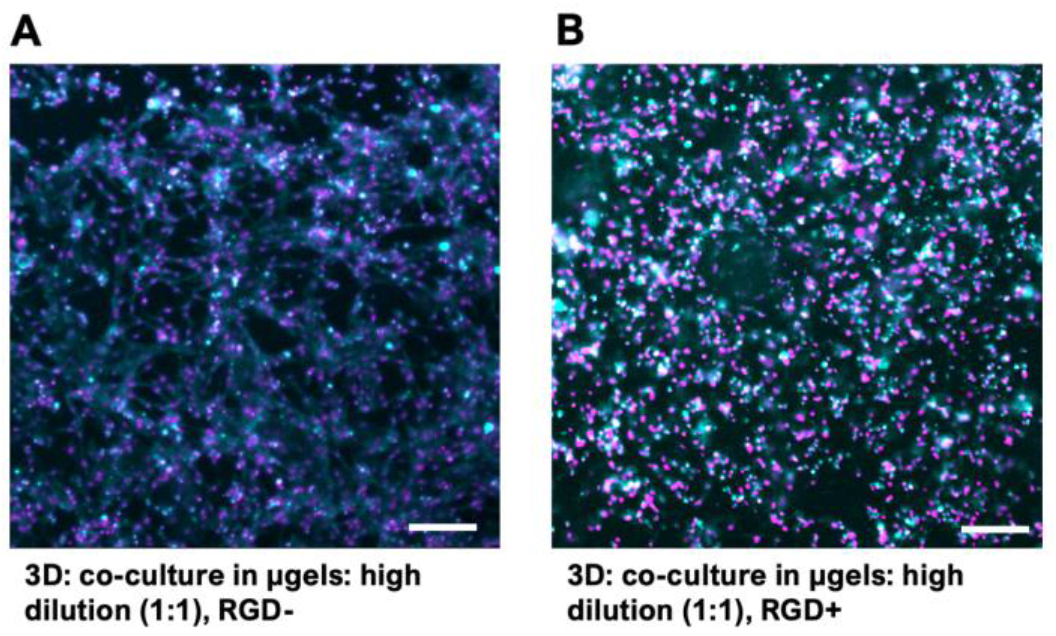
Vasculogenesis in diluted 3D granular PEG hydrogels to included cell adhesive ligand, RGD in co-culture of HUVECs with 3T3 fibroblasts. **A**. Confocal image after 24 hours exhibits microvascular-like network in the absence of RGD (RGD-). B. Confocal image after 24 hours exhibits little to no cell extension in the presence of RGD (RGD+). All scalebars = 200 μm.

To observe microvascular network formation in 3D hydrogels, we needed a positive control to confirm culture conditions for vasculogenesis, allowing us to observe morphogenesis as a function of material environment. Our PEG formulation did not include degradable crosslinks, and cells embedded within it would be entrapped and immobilized by the hydrogel network, so we looked to well-established fibrin hydrogels in which robust 3D microvascular networks form from co-cultures of HUVECs and fibroblasts^27^. Confirming our co-culture (HUVEC/fibroblast concentration and ratio) and medium formulations with growth factors were sufficient to drive vasculogenesis in a permissive 3D material environment, we observed the emergence of microvascular network structures in fibrin gels over four days. After fixation on day four, HUVECs and fibroblasts that were homogeneously distributed within the fibrin hydrogel during gel formation can be seen to have formed 3D networks (Figure 5C).

To assess whether a granular formulation of a non-degradable PEG hydrogel would permit 3D vasculogenesis, we next cultured HUVECs and fibroblasts among microgels in a granular PEG hydrogel. Using the co-culture conditions established for fibrin hydrogels^27^ with growth factor supplemented medium, we looked for the emergence of microvascular networks in 3D. Over four days in our fully jammed system, cells were observed to assemble into cell dense regions that appeared to have features of microvascular networks with respect to how cells distributed within the granular hydrogel. However, the cellular structures did not exhibit network-like connectivity of vascular branches between nodes (Figure 5D, I). We next looked to dilute the microgel system--to reduce the density of the microgels per volume with the expectation that this would both increase porous spaces and reduce physical microgel contacts, thereby reducing physical jamming forces in the granular hydrogel. Both changes to the physical environment--increased porosity and reduced jamming--should enhance cells’ abilities to move within and displace their material surroundings. By diluting the microgel density in the granular hydrogel with either 0.75 parts medium to 1 part fully jammed microgels or 1 part medium to 1 part microgels, we observed enhanced network formation (Figure 5D, II and III). From this experiment, it was evident that, within our granular hydrogel, a material environment in which cells can establish cell-cell connectivity--through one or both of porosity and material displacement--was an important step in vessel formation. Conversely, densely jammed particles appeared to limit cell-cell interactions and rearrangements, despite exhibiting macroscale stress yielding and relaxation.

Finally, towards examining the influence of cell-hydrogel adhesive interactions on vasculogenesis, we compared the emergence of vascular network structures in the granular PEG hydrogel that either included or did not include PEG-tethered RGD ligands. RGD was either natively present ^[36–38]^ or added to other continuous hydrogel systems designed to support angiogenesis ^[9,39]^ and vasculogenesis ^[10]^, and in cases where it was added it was observed to enhance network morphogenesis. Here, using our least-jammed granular PEG hydrogel for experimentation (1:1 PEG:medium), we observed the opposite effect (Figure 6). When we removed RGD from our PEG microgels, vascular network formation was enhanced (Figure 6A) compared to when RGD was present (Figure 6B and Figure 5D, III). This result was unexpected considering previous work in continuous hydrogels. The enhancement of network morphology in the absence of RGD suggests that within a granular hydrogel, a permissive environment that allows cellular activities and multicellular self-organization is centrally important for the morphogenesis of vascular structures in 3D. This suggests that integrin binding to the matrix surrounding the cells--in this case that matrix is the surfaces of PEG microgels—inhibits or slows vascular morphogenesis in this system, suggesting that either competitive interactions for cells that would otherwise be involved in vessel morphogenesis or cell signaling pathways that are induced by integrin engagement of RGD interfere or detract from vessel formation in this system.

These results open directions for further study. In 3D biomaterials, we see evidence for differential contributions of ligand binding, cell-cell interactions, material viscoelasticity, and permissivity (yielding and/or degradation) depending on material design. The specific microvascular growth process under consideration may also influence design. For example, in contrast to our results, RGD might be necessary for microvascular growth in angiogenic processes where cells are not embedded within a material but instead infiltrate from vessels surrounding an avascular volume. It is possible also that matrix deposition by fibroblasts in a coculture system may be providing cues, and the permissive nature of a granular PEG hydrogel permits its deposition and spread, which endothelial cells respond to in turn. The importance of matrixmetalloprotinase ^[10]^ activity, crucial to vasculogenesis in other 3D hydrogels, may be less important in a granular material. In a granular material, the reduction of jamming concentration might have multiple effects: both allowing space, but also reducing particle-particle contacts and physical interactions that hold microgels in place in jammed granular systems. The reduction of physical interactions may facilitate cell displacement of particles by cells ^[40]^. Determining the interplay of these factors in vascular morphogenesis will be the subject of future study.

## III. Conclusions

In this report, we have demonstrated a PEG-based granular hydrogel that supports the formation of 3D microvascular networks. This work shows that the emergence of microvascular networks from HUVECs and fibroblasts embedded within a 3D hydrogel biomaterial is possible among granular biomaterials composed of microgels with covalent intraparticle crosslinking but no particle-to-particle crosslinking. Microvascular network formation is enhanced by decreasing jamming and removing integrin binding ligands from the hydrogel surroundings. The cell-scale (<40 μm diameter) PEG microgels used here might be further engineered towards various applications through biofunctionalization, changes in polymer concentration or intraparticle crosslink design, and through interparticle crosslinking. We show that microvascular network formation can occur rapidly within a permissive environment. These observations may have implications for understanding how microvasculature develops and can be engineered into 3D biomaterials. This work will also inform approaches to vascularized granular materials, which are emerging as exciting platforms for *in vitro* and *in vivo* studies in tissue engineering and regenerative medicine.

## IV. Materials and Methods

### I. Fabrication of PEG microgels

PEG microgels were fabricated using a bulk emulsification method with a water-in-oil process. 6% (w/v) 4-arm PEG-thiol (JenKem) and 4% (w/v) 4 arm PEG-acrylate (JenKem) solutions were mixed using Dulbecco’s phosphate-buffered saline (DPBS). For experiments that incorporated the arginine-glycine-aspartic acid (RGD) cell-adhesive ligand, 1 mg/mL thiolated RGD peptide (Arg-Gly-Asp) was included in the PEG hydrogel precursor solution to facilitate cell adhesion to the surface of the microgels (Figure 1B). For fluorescence imaging of microgels, 1% (w/v) 2 MDa fluorescein isothiocyanate-tagged dextran (reagents purchest from Sigma Aldrich unless otherwise indicated) was included in the hydrogel precursor solution.

Light mineral oil including 2% (w/w) Span 80, and 0.5% triethanolamine (TEOA) was prepared to form the continuous phase of the emulsion. The mineral oil solution was placed on a stand and stirred at 2,000 rpm using an overhead stirrer. The aqueous PEG solution was placed in a 1 mL syringe with a 30-gauge needle on a syringe pump. The aqueous phase was pumped at a rate of 40 μL/minute directly into the stirred continuous phase to create a dispersion of aqueous droplets in the mineral oil. After the aqueous phase was fully dispensed into the oil, the water-in-oil suspension was allowed to stir for 45 minutes to allow the precursor solution to crosslink within the droplets.

Microgels were then recovered by transferring the suspension to a conical tube and centrifuging at 120 rcf for 5 minutes. Excess oil was removed leaving behind a pellet of microgels. To wash the microgels, isopropanol was added to the conical tube and pipetted multiple times to mix, then vortexed vigorously. The conical tube was then centrifuged again at 120 rcf for 5 minutes to pellet the microgels, and supernatant of isopropanol with residual oil was removed. Isopropanol was again added to the conical tube and then vortexed. Microgels were then separated from the isopropanol by vacuum filtration over a 0.22 μm hydrophilic polyvinylidene fluoride (PDVF) membrane. Two more washes were performed by adding isopropanol to the microgels on the membrane to completely remove excess surfactant. Finally, microgels were rehydrated with deionized water. At this point they were stored until use in forming granular hydrogels.

### II. Granular Hydrogel Formation

After filtration and cleaning, microgels were transferred to deionized water in microcentrifuge tubes. Microgels were centrifuged at 200 rcf then mixed with endothelial cell medium and placed in the fridge to be used for cell studies. Granular hydrogels were formed by centrifuging the microgels in microcentrifuge tubes at 200 rcf for 3 minutes. The endothelial cell medium supernatant was aspirated off the pellet to leave behind the fully jammed granular hydrogel. Endothelial cell medium was added at desired volumes to fully jammed granular hydrogels to create diluted granular hydrogels. The dilution ratios of granular hydrogels:cell medium used here, which changed the pore spacing between individual microgels, were 1:0 to 1:1 with 0.25 increments. Fully jammed granular hydrogels (1:0) were thus formed from microgels centrifuged at 200 rcf with no additional cell medium added in between each individual microgel. These particles were used for rheological experiments to test for granular hydrogel mechanical properties. When equal volumes of microgels and endothelial cell medium were added in a microcentrifuge tube this gave a ratio of 1:1, resulting in a granular hydrogel with more porosity.

Granular hydrogels with interparticle crosslinking to limit the flow of microgels were formed by adding 66 mM photoinitiator, lithium phenyl-2,4,6-trimethylbenzoylphosphinate (LAP), on top of the granular hydrogels and photocrosslinked at 10 mW/cm^2^ for 5 minutes to obtain a granular hydrogel in which particles were crosslinked to one another.

Granular hydrogel properties were characterized using rheological studies, and both uncrosslinked and crosslinked (interparticle) granular hydrogels were compared.

### III. Cell Culture

Human umbilical vein endothelial cells (HUVECs) were used for all experiments and cultured (passages 5-8) in EGM-2 bulletkit medium (Lonza). Culture medium was changed every two days. NIH 3T3 Fibroblasts were cultured (passages 10-13) in Dulbecco’s modified Eagle’s medium (DMEM/F-12) medium (Gibco) with 10% (v/v) fetal bovine serum and 1% (v/v) anti-biotic/anti-mycotic solution. Medium was changed every two days. Cells were passaged when cells reached 70-80% confluency.

For cell experiments, HUVECs and fibroblasts were mixed at a ratio of 5:1 at concentrations of 5 million cells/mL and 1 million cells/mL, respectively, and combined with PEG microgels. Co-culture experiments to generate microvascular networks were done in EGM-2 bulletkit medium supplemented with 100 ng/mL recombinant human angiopotetin-1 (Peprotech) and 50 ng/mL human VEGF-165A (ACROBiosystems).Medium in the co-culture environment was changed every two days. Cell morphology was observed after 24 hours and 4 days. Images were taken at each timepoint.

The polydimethylsiloxane (PDMS) device used to contain the granular hydrogel during experiments with cells was cast from a mold created using Autodesk Fusion360 and printed with Formlabs Form 2 3D printer. The mold was glued on a microscope glass slide. PMDS was prepared by mixing elastomer base and curing agent vigorously at a 10:1 weight ratio. PDMS solution was placed under vacuum to remove any air bubbles formed while mixing and was then poured into the mold. The mold was placed in a 37ºC incubator and left overnight for curing. Once cured, the PDMS device was removed from the mold. New slides were plasma-treated and the PDMS device was placed on them for attachment. Devices were immersed in 70% ethanol for sterilization before use in cell studies.

### IV. Formation of continuous PEG hydrogels

Continuous PEG hydrogels were fabricated with 6% (w/v) 4-arm PEG-thiol and 4% (w/v) 4 arm PEG-acrylate solutions prepared in DPBS. We pipetted 500 μL of the PEG precursor solution into each well of a 24 well plate to cover the entire bottom surface and let sit at room temperature for a few hours until gelation. We seeded cells on top of the PEG hydrogels with RGD (and without RGD as a control), centrifuged at 200 rcf for even cell distribution, and added culture medium to each well. We subsequently handled cell culture as described previously.

### V. Formation of cell-containing fibrin gels

Fibrinogen was dissolved in PBS at 20 mg/mL, twice the desired final concentration. Thrombin was dissolved in PBS at 10 U/mL. HUVECs and fibroblasts were mixed at a ratio of 5:1 and spun down at 200 rcf for 5 minutes. The cell pellet was resuspended in endothelial cell medium containing 2 U/mL of thrombin over ice. The solution was mixed with fibrinogen solution over ice at a 1:1 ratio to produce a final fibrinogen solution with a desired concentration of 10 mg/mL, then pipetted into the PDMS devices immediately.We subsequently handled cell culture as described previously.

### VI. Formation of cell-containing granular hydrogels

Microgel dilutions were achieved by centrifuging microgels alone at 3,095 rcf for 5 minutes in microcentrifuge tubes. We aspirated off the endothelial cell medium supernatant leaving fully jammed microgels behind and pipetted 100 μL microgels into a fresh microcentrifuge tube. Desired volumes of fresh endothelial cell medium mixed with cell concentrations were added to the microgels to obtain varied ratios of microgels:cell medium (e.g., 100 μL microgels with 100 μL medium = 1:1 ratio and 100 μL microgels with no medium = 1:0 ratio, which were considered fully jammed particles). We homogeneously mixed PEG microgels containing RGD with HUVECs and fibroblasts at a ratio of 5:1 in endothelial cell medium. We placed microgels and cells in the PDMS culturing device for cell culture according to the methods described above. Samples were fixed after four days to examine vessel networks.

### VII. Cell Staining

For analysis of cell experiments, HUVECs were stained with CellTracker Deep Red fluorescent dye (Invitrogen) to examine cellular movement during experimentation. Co-cultures were fixed with 4% paraformaldehyde for 15 minutes before 1 hour permeabilization and blocking with a 0.1% (v/v) Triton X-100 solution in PBS with bovine serum albumin (BSA, 3% (w/v)) to prevent nonspecific binding. Samples were washed with PBS three times for 5 minutes each. Samples were then counterstained with primary antibodies at room temperature. Primary antibodies used in this work included donkey to anti-rabbit 647 (Abcam, ab150075) as well as phalloidin 568, which labels F-actin, in blocking solution for 1 hour. Lastly, samples were washed twice with PBS for 5 minutes with DAPI, which labels nuclear DNA, added into the second wash. All samples were protected from light and stored at 4°C until imaging.

### VIII. Imaging and Image Analyses

Imaging was conducted on a Leica DMi8 widefield microscope and Olympus FV1000 confocal microscope. Images containing microgels and cells were collected from microscope slides and imaged from the bottom up. Other images containing PEG hydrogel and cells were collected on 25 × 25 mm glass coverslips. Imaging settings (exposure time and light intensity) were held constant for all imaging where fluorescence intensities were compared across multiple samples. At least three distinct areas per scaffold were imaged for cell morphology analyses. Images were processed using Fiji software.

## References

1. Duval, K. et al. Modeling physiological events in 2D vs. 3D cell culture. Physiology 32, 266–277 (2017).

2. Moon, J. J. et al. Biomimetic hydrogels with pro-angiogenic properties. (2010) doi:10.1016/j.biomaterials.2010.01.104.

3. Zhao, X. et al. Review on the Vascularization of Organoids and Organoids-on-a-Chip. Front Bioeng Biotechnol 9, 637048 (2021).

4. Koroleva, A. et al. Hydrogel-based microfluidics for vascular tissue engineering. BioNanoMaterials 17, 19–32 (2016).

5. Cuchiara, M. P., Gould, D. J., Mchale, M. K., Dickinson, M. E. & West, J. L. Integration of Self-Assembled Microvascular Networks with Microfabricated PEG-Based Hydrogels. (2012) doi:10.1002/adfm.201200976.

6. Kolesky, D. B., Homan, K. A., Skylar-Scott, M. A. & Lewis, J. A. Three-dimensional bioprinting of thick vascularized tissues. Proc Natl Acad Sci U S A 113, 3179–3184 (2016).

7. Lin, C. C. & Anseth, K. S. PEG hydrogels for the controlled release of biomolecules in regenerative medicine. Pharm Res 26, 631–643 (2009).

8. Peters, E. B., Christoforou, N., Leong, K. W., Truskey, G. A. & West, J. L. Poly(Ethylene Glycol) Hydrogel Scaffolds Containing Cell-Adhesive and Protease-Sensitive Peptides Support Microvessel Formation by Endothelial Progenitor Cells. Cell Mol Bioeng 9, 38–54 (2016).

9. Song, K. H., Highley, C. B., Rouff, A. & Burdick, J. A. Complex 3D-Printed Microchannels within Cell-Degradable Hydrogels. Adv Funct Mater 28, 1801331 (2018).

10. Wei, Z., Schnellmann, R., Pruitt, H. C. & Gerecht, S. Hydrogel Network Dynamics Regulate Vascular Morphogenesis. Cell Stem Cell 27, 798-812.e6 (2020).

11. Highley, C. B., Song, K. H., Daly, A. C. & Burdick, J. A. Jammed Microgel Inks for 3D Printing Applications. Advanced Science 6, (2019).

12. Miller, J. S. et al. Rapid casting of patterned vascular networks for perfusable engineered 3D tissues. Nat Mater 11, 768 (2012).

13. Menut, P., Seiffert, S., Sprakel, J. & Weitz, D. A. Does size matter? Elasticity of compressed suspensions of colloidal- and granular-scale microgels. Soft Matter 8, 156–164 (2011).

14. Daly, A. C., Riley, L., Segura, T. & Burdick, J. A. Hydrogel microparticles for biomedical applications. Nature Reviews Materials 2019 5:1 5, 20–43 (2019).

15. Griffin, D. R., Patterson, J. T. & Kasko, A. M. Photodegradation as a mechanism for controlled drug delivery. Biotechnol Bioeng 107, 1012–1019 (2010).

16. Bhattacharjee, T. et al. Liquid-like Solids Support Cells in 3D. ACS Biomater Sci Eng 2, 1787–1795 (2016).

17. Shiwarski, D. J., Hudson, A. R., Tashman, J. W. & Feinberg, A. W. Emergence of FRESH 3D printing as a platform for advanced tissue biofabrication. APL Bioeng 5, 010904 (2021).

18. Highley, C. B., Rodell, C. B. & Burdick, J. A. Direct 3D Printing of Shear-Thinning Hydrogels into Self-Healing Hydrogels. Advanced Materials 27, 5075–5079 (2015).

19. Griffin, D. R., Weaver, W. M., Scumpia, P. O., di Carlo, D. & Segura, T. Accelerated wound healing by injectable microporous gel scaffolds assembled from annealed building blocks. Nat Mater 14, 737–744 (2015).

20. Bhattacharjee, T. et al. Writing in the granular gel medium. Sci Adv 1, (2015).

21. Chen, M. H. et al. Injectable Supramolecular Hydrogel/Microgel Composites for Therapeutic Delivery. Macromol Biosci 19, 1800248 (2019).

22. Muir, V. G., Qazi, T. H., Shan, J., Groll, J. & Burdick, J. A. Influence of Microgel Fabrication Technique on Granular Hydrogel Properties. ACS Biomater Sci Eng 7, 4269–4281 (2021).

23. Chaudhuri, O. et al. Substrate stress relaxation regulates cell spreading. (2015) doi:10.1038/ncomms7365.

24. Sessoms, D. A., Bischofberger, I., Cipelletti, L. & Trappe, V. Multiple dynamic regimes in concentrated microgel systems. Philosophical Transactions of the Royal Society A: Mathematical, Physical and Engineering Sciences 367, 5013–5032 (2009).

25. Broguiere, N. et al. Morphogenesis Guided by 3D Patterning of Growth Factors in Biological Matrices. Advanced Materials 32, 1908299 (2020).

26. Bhattacharjee, T. & Angelini, T. E. 3D T cell motility in jammed microgels. J Phys D Appl Phys 52, 024006 (2019).

27. Whisler, J. A., Chen, M. B. & Kamm, R. D. Control of Perfusable Microvascular Network Morphology Using a Multiculture Microfluidic System. doi:10.1089/ten.tec.2013.0370.

28. Chaudhuri, O. et al. Hydrogels with tunable stress relaxation regulate stem cell fate and activity. Nat Mater 15, 326–334 (2016).

29. Cuchiara, M. P., Gould, D. J., McHale, M. K., Dickinson, M. E. & West, J. L. Integration of Self-Assembled Microvascular Networks with Microfabricated PEG-Based Hydrogels. Adv Funct Mater 22, 4511–4518 (2012).

30. Chapla, R. & West, J. L. Hydrogel biomaterials to support and guide vascularization. Progress in Biomedical Engineering 3, 012002 (2021).

31. Brassard, J. A. & Lutolf, M. P. Engineering Stem Cell Self-organization to Build Better Organoids. Cell Stem Cell 24, 860–876 (2019).

32. Douglas, A. M. et al. Dynamic assembly of ultrasoft colloidal networks enables cell invasion within restrictive fibrillar polymers. Proc Natl Acad Sci U S A 114, 885–890 (2017).

33. Ford, M. C. et al. A macroporous hydrogel for the coculture of neural progenitor and endothelial cells to form functional vascular networks in vivo. Proc Natl Acad Sci U S A 103, 2512–2517 (2006).

34. de Rutte, J. M., Koh, J. & di Carlo, D. Scalable High-Throughput Production of Modular Microgels for In Situ Assembly of Microporous Tissue Scaffolds. Adv Funct Mater 29, (2019).

35. Seymour, A. J., Shin, S. & Heilshorn, S. C. 3D Printing of Microgel Scaffolds with Tunable Void Fraction to Promote Cell Infiltration. Adv Healthc Mater 10, 2100644 (2021).

36. Jeon, J. S. et al. Generation of 3D functional microvascular networks with human mesenchymal stem cells in microfluidic systems. Integrative Biology (United Kingdom) 6, 555–563 (2014).

37. Nguyen, D. H. T. et al. Biomimetic model to reconstitute angiogenic sprouting morphogenesis in vitro. Proc Natl Acad Sci U S A 110, 6712–6717 (2013).

38. Moya, M. L., Hsu, Y. H., Lee, A. P., Christopher, C. W. H. & George, S. C. In vitro perfused human capillary networks. Tissue Eng Part C Methods 19, 730–737 (2013).

39. Leslie-Barbick, J. E., Saik, J. E., Gould, D. J., Dickinson, M. E. & West, J. L. The promotion of microvasculature formation in poly(ethylene glycol) diacrylate hydrogels by an immobilized VEGF-mimetic peptide. Biomaterials 32, 5782–5789 (2011).

40. Douglas, A. M. et al. Dynamic assembly of ultrasoft colloidal networks enables cell invasion within restrictive fibrillar polymers. Proc Natl Acad Sci U S A 114, 885–890 (2017).

